# Endothelial ANGPT2 insufficiency impairs retinal vascularization with ROP-like neovascular tufts

**DOI:** 10.64898/2026.08.14.744858

**Authors:** Zhiliang Sun, Kai Ding, Taotao Li, Jinting Zhang, Xin Shen, Xiwen Jia, Xiao Li, Xudong Cao, Beibei Xu, Peirong Lu, Yulong He

## Abstract

ANGPT2 is widely recognized as a critical regulator of pathological neovascularization. By analyzing scRNA-seq data from neonatal retinas, we demonstrate that *Angpt2* transcripts are highly enriched in tip cells relative to other endothelial subtypes, where *Angpt1*/*4* expression is absent. However, mechanisms underlying ANGPT2 function at angiogenic fronts remain inadequately understood. Here, we show that endothelial *Angpt2* deletion severely disrupted retinal vascularization, characterized by neovascular tufts and micro-hemorrhage. Similar angiogenic defects also occurred in the brain, but were less evident in other tissues examined. Mechanistically, ANGPT2 insufficiency attenuated retinal tip cell invasion with aberrant mural cell coverage, compromising sprouting into non-vascularized tissues. Retinal RNA-seq analysis revealed that transcripts associated with endothelial migration and junction assembly were reduced in *Angpt2* mutants compared to littermate controls, while upregulated genes were enriched in hypoxia-responsive pathways and mural cell development. Notably, abnormal mural-tip cell associations were detected within 48 hours post-*Angpt2* deletion, displaying also a hypoxia-driven transcriptomic signature. These closely resemble the vascular pathologies observed in human retinopathy of prematurity. In contrast, *Angpt1* insufficiency or *Angpt4* deficiency primarily affected venous morphogenesis. Collectively, our findings imply that ANGPT2 is essential for driving tip cell invasion during sprouting angiogenesis, and that its insufficiency triggers hypoxia-driven vascular anomalies.

## Introduction

During early retinal vascularization, specialized endothelial cells (ECs) known as “tip cells” lead the vascular front. Guided by angiogenic growth cues, tip cells migrate toward the non-vascularized hypoxic regions, while trailing stalk cells proliferate to elongate the new vessels for establishing a vascular network [1, 2]. The identity of tip ECs is transient and appears to be regulated by angiogenic factors including VEGFA-mediated signaling pathway [3, 4]. ESM1 (endothelial cell-specific molecule 1) was shown to modulate tip EC behavior via the modulation of VEGFA availability [5]. A study by Pitulescu et al. further demonstrated that DLL4 expression by tip ECs is not required to maintain tip cell identity or prevent replacement by neighboring cells. Instead, DLL4-NOTCH signaling acts as a critical driver for steering tip-derived cells into developing arteries [6]. Alongside angiogenic sprouting, mural cell recruitment to newly formed vessels is crucial for ensuring vessel stability during the establishment of a functional vascular system [7, 8]. Particularly, pericyte coverage is crucial for the blood-retinal barrier [9]. The EC-derived PDGFB (platelet-derived growth factor-B) acts as a chemoattractant to drive mural cell migration and proliferation along the new vessel [10, 11]. TGFβ promotes the differentiation of mesenchymal progenitor cells into mature mural cells [12]. Netrin-4 and Nitric Oxide (NO) have been shown to promote vascular muscle cell recruitment or the directional migration of mural cell precursors toward ECs [13]. The mural-EC interaction requires NOTCH3 signaling for mural cell maturation to promote vessel patterning [7, 14].

The angiopoietin (ANGPT)-TIE receptor pathway mediates signals for EC survival and vascular stability [15, 16]. ANGPT1, which is produced by mural cells and platelets [17, 18], exerts its primary functions through the TIE2 receptor; this includes promoting the assembly of EC cell-cell junctions [19, 20]. Additionally, the ANGPT1-TIE2 pathway is crucial for establishing venous endothelial identity, while TIE1 acts synergistically with TIE2 to restrict sprout angiogenesis during venous assembly [21–23]. Similarly, loss of ANGPT4 has been shown to impair venous maturation in the retina [24]. In contrast, ANGPT2 regulates angiogenesis by a context-dependent interaction with TIE2 and also induces integrin-mediated RAC1 activation for EC migration [25, 26]. While TIE2 and ANGPT1 are weakly expressed or undetectable in tip ECs, ANGPT2 is highly expressed in these cells and upregulated by hypoxia [22, 27, 28].

Retinopathy of prematurity (ROP) is a biphasic, hypoxia-driven vasoproliferative disorder that remains a leading cause of childhood blindness worldwide. The condition is characterized by initial vessel cessation followed by abnormal, hyperpermeable angiogenesis, culminating in the formation of pathological neovascular tufts. While the roles of VEGF in inducing vascular permeability and endothelial proliferation are well-documented, the local regulators involved in ROP progression at angiogenic fronts remain inadequately understood. Here, we demonstrated that ANGPT2 insufficiency replicated the core pathological hallmarks of ROP. Specifically, endothelial *Angpt2* deletion retarded endothelial tip cell migration and shifted the molecular landscape toward a hypoxia-driven transcriptional signature. This genetic insufficiency disrupts the retinal vascularization, manifesting as aberrant mural cell recruitment around tip cells, a massive expansion of ESM1-positive endothelial cells, compromised vascular integrity, and the formation of neovascular tufts. By showing that ANGPT2 insufficiency triggers these hypoxia-driven vascular anomalies, our findings provide new insights into the molecular mechanisms driving ROP-like pathologies.

## Results

### Neovascular tuft formation upon endothelial *Angpt2* deletion

To investigate the role of Angiopoietin-2 (ANGPT2) across different endothelial cell (EC) subtypes, we re-analyzed published single-cell RNA sequencing data from the postnatal day 6 (P6) mouse retina [27]. Based on marker gene expression, retinal ECs clustered into five distinct populations (Fig. 1A). Notably, *Angpt2* was highly enriched in the tip EC cluster, whereas its expression was substantially lower in the other EC subtypes. Interestingly, *Angpt1* and *Tek* expression was low or undetectable in tip cells (Fig. 1B-C). Given that ANGPT2 is predominantly expressed in endothelial cells, particularly tip ECs, we generated an endothelial-specific inducible knockout mouse model (*Angpt2^Flox/−^*;*Cdh5-Cre^ERT2^*, hereafter referred to as *Angpt2^iKO;CDH5+^*), using *Angpt2^Flox/+^*;*Cdh5-Cre^ERT2^* littermates served as controls. Following tamoxifen treatment from postnatal days 1 to 4 (P1 to P4), we analyzed ANGPT2 protein expression and retinal vascularization. At P7, a reduction of approximately 70% in ANGPT2 protein levels was detected in the lung tissue of *Angpt2^iKO;CDH5+^* mice compared to controls (Fig. 1D-F); the residual expression was likely attributable to the remaining heterozygous endothelial cells or non-endothelial cells. Immunofluorescence staining revealed impaired retinal vascularization in *Angpt2^iKO;CDH5+^* mice from P7 to P21, characterized by a significant decrease in the vascularization index, defined as the ratio of vascularized area to total retinal area (Fig. 1G-H). Strikingly, vein-associated tufts-like structures emerged by P15 and became more pronounced by P21, indicating that postnatal endothelial-ANGPT2 insufficiency drives the formation of neovascular tufts (Fig. 1H-I).

**Fig. 1.**
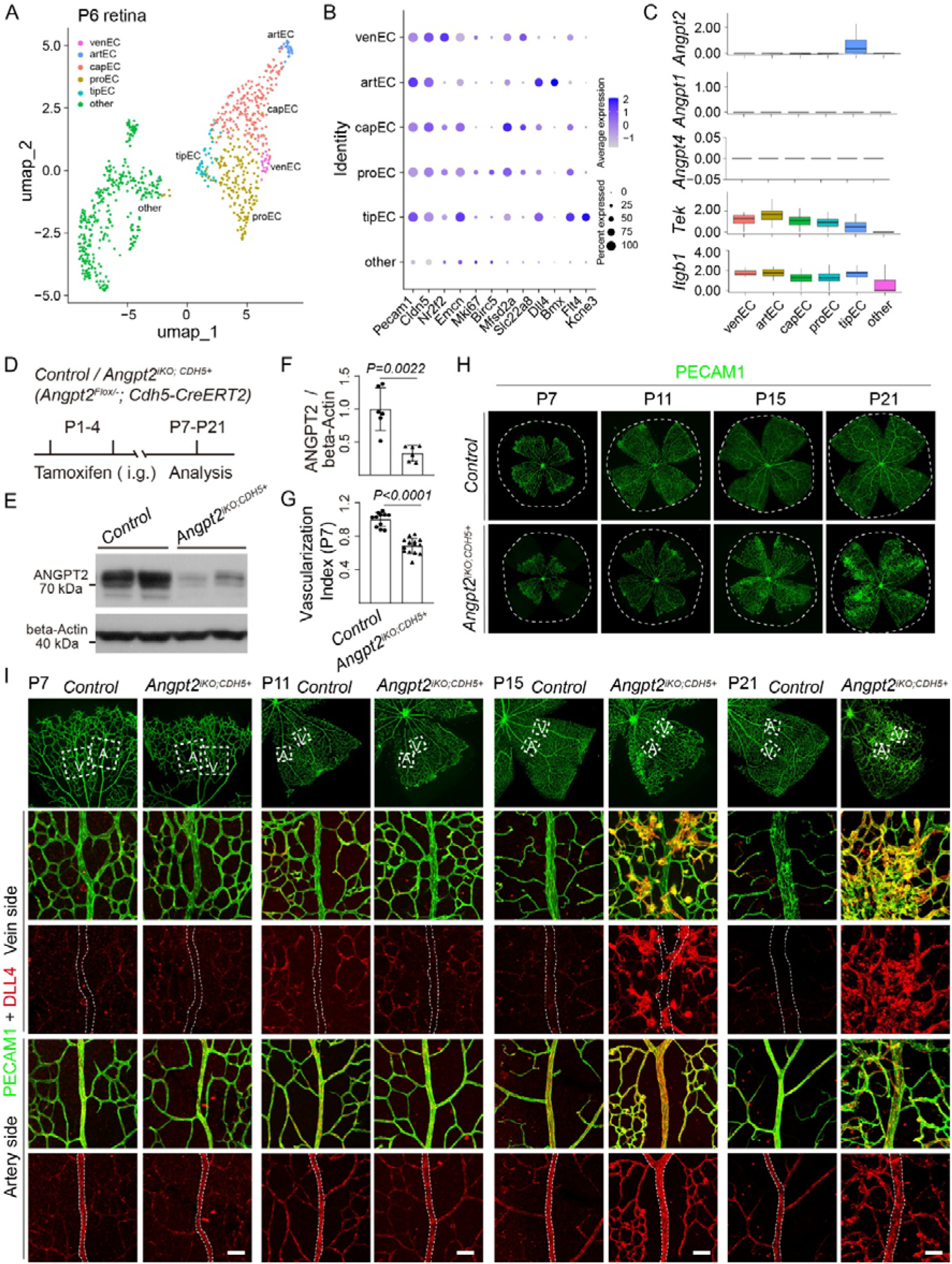
Analysis of ANGPT2 expression in tip ECs by the scRNA-seq and its function in the retinal vascular morphogenesis. **A-C**. Single-cell RNA-seq analysis of endothelial cells (ECs) in mouse retinas at the postnatal day 6 (P6). UMAP plot shows the EC clustered into five major subtypes (A), including venEC (venous EC), artEC (artery EC), capEC (capillary EC), tipEC (tip EC), proEC (proliferated EC) according to the marker genes (B). *Angpt2* expression is markedly enriched in tipECs compared with other EC subtypes, where *Angpt1* or *Angpt4* is minimal or absent (C). *Itgb1* is highly expressed in tip ECs while *Tek* is low in tip cells relative to other EC subtypes. **D**. Tamoxifen intragastric (i.g.) administration and the analysis scheme. **E**. Deletion efficiency of ANGPT2 was assessed by the Western blotting analysis of lung tissues from *Angpt2^iKO;CDH5+^*and control mice at P7. **F**. Quantification of ANGPT2 protein normalized to beta-actin (ANGPT2/beta-actin: Control: 1.00±0.32, n=6; *Angpt2^iKO;CDH5+^*: 0.33±0.12, n=6, *P*=0.0022). **G**-**H**, Retinal blood vessel analysis of *Angpt2^iKO;CDH5+^* and control mice between P7 and P21 (Quantification of the vascularization index, defined as the ratio of vascularized area to total retinal area, normalized to littermate controls at P7; Control: 1.00±0.08, n=11; *Angpt2^iKO;CDH5+^*: 0.69±0.01, n=15, *P*<0.0001). **I**. Visualization of retinal blood vessels of *Angpt2^iKO;CDH5+^* mice by immunostaining at P7, P11, P15 and P21. Scale bar: 50 μm in I.

To further validate the role of ANGPT2 in retinal vascular development, we generated another inducible *Angpt2* knockout mouse model using the *Ubc-Cre^ERT2^* strain (*Angpt2^Flox/−^*;*Ubc-Cre^ERT2^*, hereafter designated as *Angpt2^iKO;UBC+^*). *Angpt2^Flox/+^*;*Ubc-Cre^ERT2^*littermates was utilized as controls. Following tamoxifen administration from postnatal P1 to P4, successful deletion of *Angpt2* was confirmed in lung tissues at P7 (Supplemental Fig. 1A-C). In agreement with prior reports, retinal vascular analysis at P7 revealed a marked reduction in both the vascularized area and capillary density within the regions between mature arterioles and venules in *Angpt2^iKO;UBC+^* mice compared to littermate controls (Supplemental Fig. 1D). Quantitatively, the vascularization index demonstrated an approximately 30% reduction in *Angpt2^iKO;UBC+^* mice (Supplemental Fig. 1E).

In addition, to specifically investigate the role of ANGPT2 expressed by hematopoietic cells, such as macrophages, we generated a hematopoietic cell-specific *Angpt2* knockout model by crossing *Angpt2^Flox/Flox^* mice with the *Vav-iCre* strain (*Angpt2^Flox/Flox^*; *Vav-iCre*, hereafter designated as *Angpt2^KO;VAV+(Vav-iCre)^*). *Angpt2^Flox/Flox^* littermates served as controls. Notably, immunofluorescence staining demonstrated that retinal vascular development was unaffected in *Angpt2^KO;VAV+(Vav-iCre)^* mice at P21 (Supplemental Fig. 1F).

### Defective tip EC migration during retinal angiogenesis

To assess the structural consequences of endothelial *Angpt2* deletion induced from P1 to P4, cryosections of P15 mouse eye tissue were analyzed. The results confirmed underdevelopment or absence of the vascular networks in the deep and intermediate retinal layers of *Angpt2^iKO;CDH5+^* mice, along with delayed hyaloid vessel degeneration (Fig. 2A-B). At P21, *Angpt2^iKO;CDH5+^* retinas exhibited a loss of vessels in the third layer on the venous side and an incomplete vascular network in the second layer on the arterial side, further indicating that endothelial ANGPT2 deficiency impairs the migration of retinal endothelial cells into the deeper tissue layers (Fig. 2C). Immunofluorescence staining for the mural cell marker NG2 revealed abnormal pericyte recruitment on vascular tufts in P21 *Angpt2^iKO;CDH5+^* retinas (Fig. 2D). Consistently, aberrant mural cell coverage was already detectable on the retinal tip cells of P7 *Angpt2^iKO;CDH5+^*mice (Fig. 2E). Given the role of pericytes in regulating angiogenesis and maintaining vascular stabilization, we hypothesize that the tuft-like vascular structures observed in the retinas with endothelial *Angpt2* deletion result from impaired endothelial cell migration into deeper retinal layers.

**Fig. 2.**
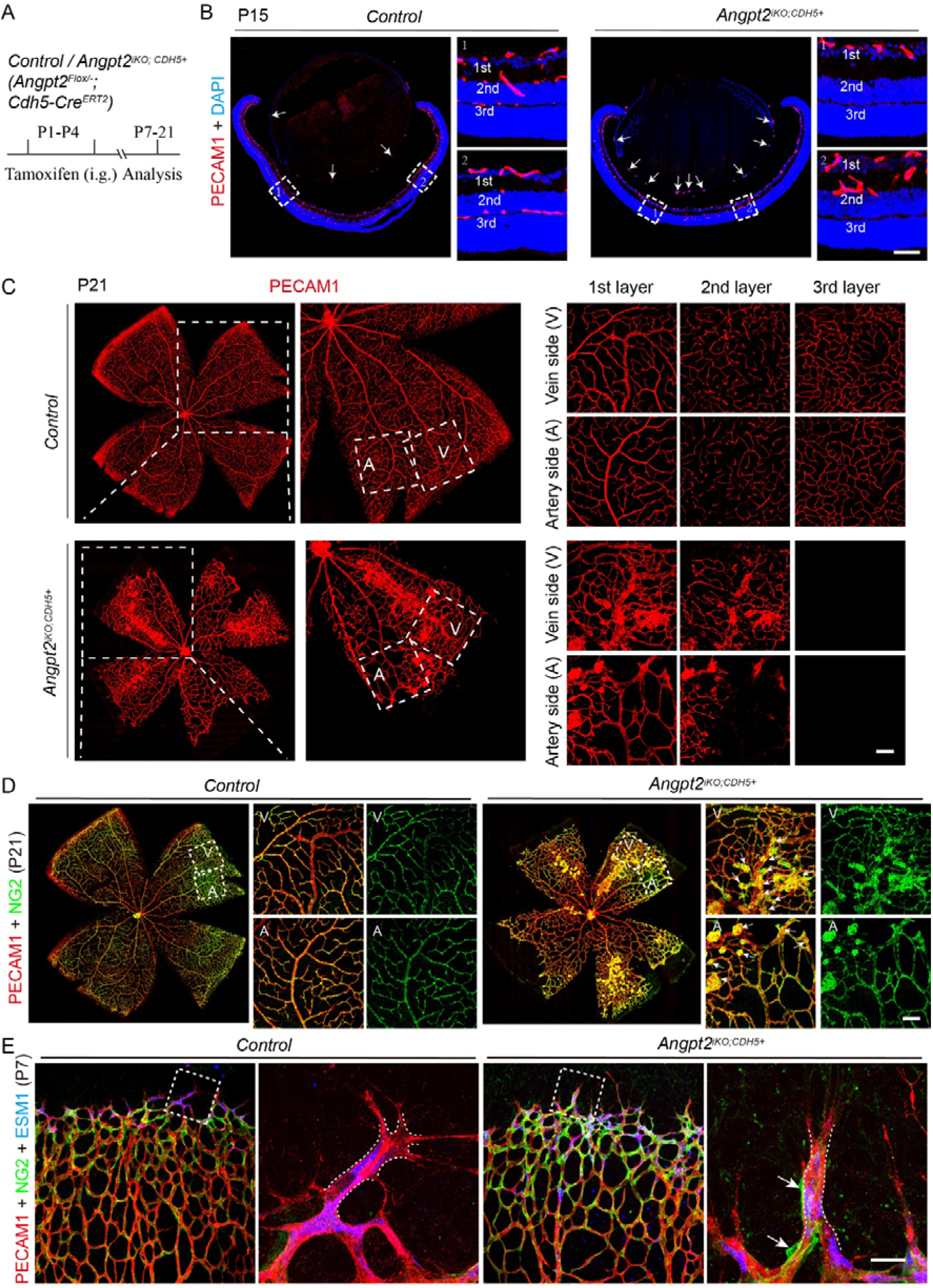
Retardation of the retinal tip EC migration with aberrant mural cell coupling after the induced deletion of endothelial *Angpt2*. **A.** Tamoxifen intragastric (i.g.) administration and analysis scheme. B. Cross-sectional analysis of the three layers of retinal blood vessels in *Angpt2* mutant and littermate control mice at P15. Arrows point to hyaloid vessels. **C**. Analysis of the three layers of retinal blood vessels in *Angpt2^iKO;CDH5+^* and control mice by whole-mount immunostaining at P21. **D** Analysis of pericytes coverage of retinal tip ECs in *Angpt2^iKO;CDH5+^* and control mice by immunostaining at P21. Arrows indicate vascular tufts. **E**. Immunostaining for NG2 in retinas from *Angpt2*-deleted and littermate control mice at P7. Arrows highlight pericyte abnormal recruitment in tip cells. Scale bar: 50 μm in B; 100 μm in C and D; 20 μm in E.

### Induction of a hypoxia-driven transcriptomic signature upon ANGPT2 insufficiency

To further investigate the molecular mechanisms by which ANGPT2 regulates retinal angiogenesis, transcriptome sequencing was performed on retinal tissue from P7 *Angpt2^iKO;CDH5+^* mice and littermate controls. Volcano plots and heatmaps displayed the differentially expressed genes (DEGs) revealed that many upregulated genes were associated with retinal hypoxia, whereas downregulated genes were linked to retinal vessel formation and integrity (Fig. 3A-C). Specifically, the hypoxia-related genes such as *Vegfa*, *Apln*, *Esm1*, and *Dll4* were significantly upregulated, while *Kdr* and *Pdgfb* exhibited an upward trend. In contrast, the vascular integrity gene *Tek* was significantly downregulated, and *Tie1*, *Cldn5*, and *Cdh5* showed a downward trend (Fig. 3D). Gene set enrichment analysis (GSEA) demonstrated significant enrichment of biological processes related to endothelial cell migration, tip and structural ECs, angiogenesis, and hypoxia (Fig. 3E). Although previous studies indicate that elevated ANGPT2 disrupts endothelial stability and increases vascular permeability, we unexpectedly observed TER119^+^ erythrocytes leaking into the perivascular space of P21 *Angpt2^iKO;CDH5+^* mice retinas, primarily accumulating adjacent to capillaries (Fig. 3F). Immunofluorescence staining for the endothelial junction proteins CLDN5 and CDH5 revealed the defective junctional organization, confirming compromised vascular integrity (Fig. 3F). GO analysis showed that downregulated genes were enriched in retinal vascular morphogenesis and integrity, whereas upregulated genes clustered in pathways associated with hypoxic responses and vascular smooth muscle cell differentiation and proliferation (Fig. 3G-H). Together, these transcriptomic findings are highly consistent with the observed retinal vascular phenotypes.

**Fig. 3.**
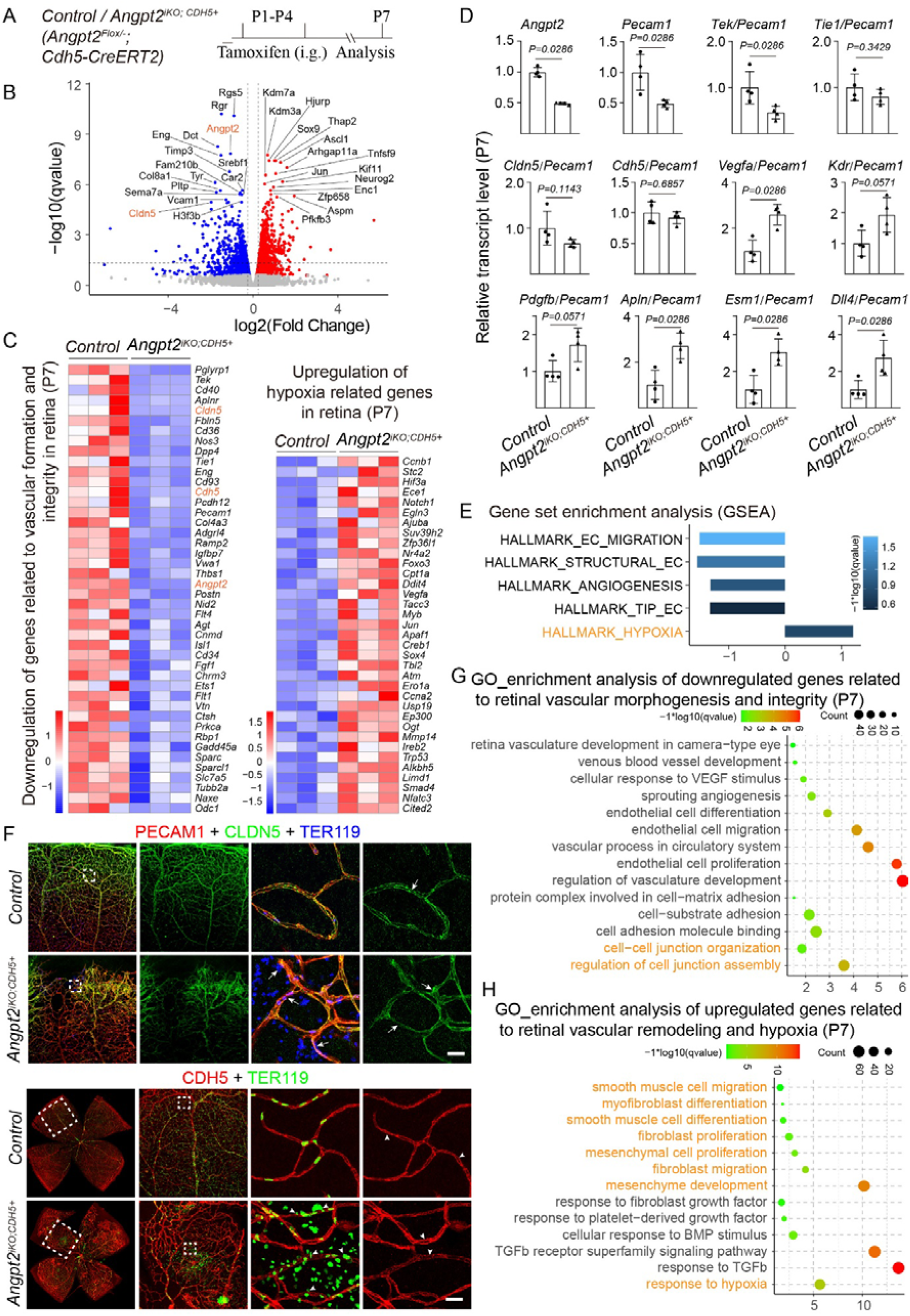
ANGPT2 insufficiency induces hypoxia-driven vascular defects in retinas. **A.** Tamoxifen intragastric (i.g.) administration and analysis scheme. **B**-**D**, RNA-seq analysis of the retinal tissues from *Angpt2* mutant (*Angpt2^iKO;CDH5+^*) and control mice at P7. Volcano plot showing differentially regulated genes (B). Heatmaps displaying subsets of upregulated hypoxia-related genes and downregulated vascular formation and integrity genes (*P*<0.05, C). Quantitative expression analysis of vascular integrity genes (*Tek*, *Tie1*, *Cldn5*, *Cdh5*) and hypoxia-associated genes (*Vegfa*, *Kdr*, *Pdgfb*, *Apln*, *Esm1*, *Dll4*) in the retina of *Angpt2^iKO;CDH5+^* and littermate control mice at P7 (D). Gene Set Enrichment Analysis (GSEA) revealed significant downregulation of gene sets related to endothelial cell migration, endothelial cell structure, angiogenesis, and tip endothelial cells following *Angpt2* deletion, accompanied by marked upregulation of hypoxia-related genes (E). **F**. Immunostaining of retinas at P21. PECAM1, CLDN5 and TER119 staining in *Angpt2* deleted and littermate control mice. Arrowheads indicate abnormal CLDN5 expression. (F) CDH5 and TER119 staining in *Angpt2* mutant and control mice. Arrows indicate abnormal CDH5 expression. **G-H**. Gene Ontology (GO) enrichment analysis. Downregulated genes were enriched for pathways related to retinal vessels formation and integrity (G). Upregulated genes were enriched for pathways associated with hypoxia and the mural cell related vascular remodeling (H). Scale bar: 20 μm in F.

### Conservation of ANGPT2 function in cerebrovascular angiogenesis

Because retinal and cerebral vessels are both part of the central nervous system vasculature and share structural and functional similarities, cerebral vessel development was also examined at P7 and P21 following endothelial-induced deletion from P1 to P4. Similar to the phenotype observed in the retina, the brains of P21 *Angpt2^iKO;CDH5+^* mice exhibited impaired angiogenesis characterized by tuft-like vascular structures and hemorrhagic pathology, although no significant bleeding was observed at P7 (Fig. 4A-C). Additionally, we assessed the effects of postnatal endothelial-ANGPT2 insufficiency on the vascular networks of the skin and intestinal villi. No significant differences in vascular density or integrity were observed in the skin of *Angpt2^iKO;CDH5+^*mice compared with littermate controls at P7 (Supplemental Fig. 2A-B). In contrast, the intestinal villi, characterized by high regeneration and metabolic activity, showed a significant reduction in vascular network area at P21, although vascular integrity appeared intact (Supplemental Fig. 2C-D).

**Fig. 4.**
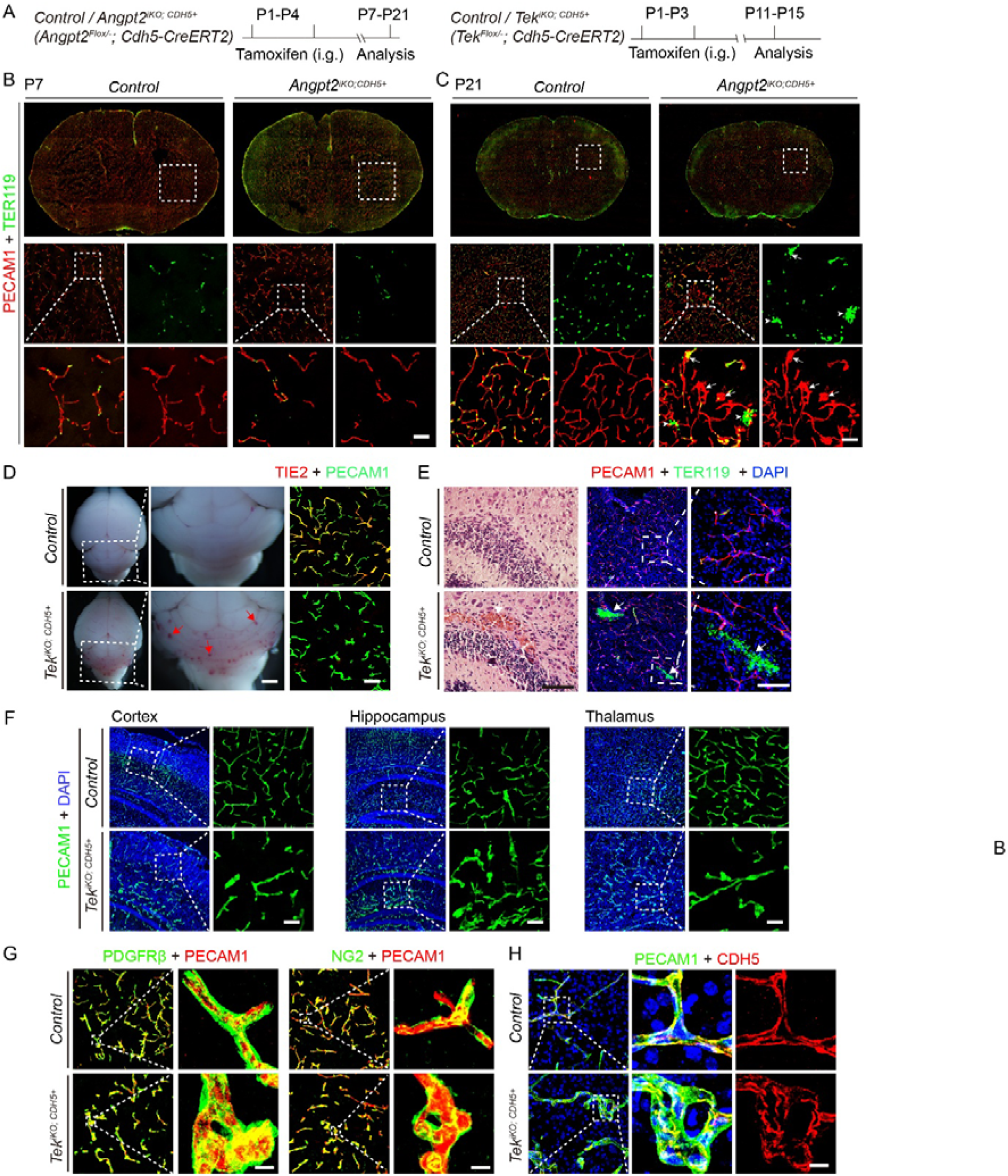
Phenotypic comparison of endothelial *Angpt2* and *Tek* deletion in cerebrovascular development. **A**. Tamoxifen intragastric (i.g.) administration and analysis scheme. **B**. Immunostaining for PECAM1 (red) and Ter119 (green) in brain of *Angpt2^iKO;CDH5+^* and littermate control mice at P7. **C**. Evidence of bleeding and neovascular tufts in brain blood vessels of *Angpt2^iKO;CDH5+^* mice at P21. Arrows indicate vascular tufts, arrowheads indicate red blood cells outside of blood vessels. **D.** Left: Stereomicroscopy images of whole brains; red arrows point to vascular hemorrhage in the hindbrain. Right: Co-immunostaining for PECAM1 (green) and TIE2 (red) in cerebral blood vessels at P11 to evaluate *Tek* deletion efficiency. **E.** Left: H&E staining of the cerebellum. Right: Immunostaining of coronal brain sections for PECAM1 (red), TER119 (green), and DAPI (blue); white arrows point to areas of hemorrhage. **F.** Immunostaining of coronal brain sections for PECAM1 (green) and DAPI (blue) across multiple regions, including the cortex, hippocampus, and thalamus at P15. **G.** Immunostaining of coronal brain sections for PECAM1 (red) counterstained with either PDGFRβ (green) or NG2 (green). H. Immunostaining of coronal brain sections for PECAM1 (green) and CDH5 (red). Scale bar: 50 μm in B, C and F; 1 mm in D (left panel); 100 μm in D (right panel) and E; 10 μm in G and H.

Because ANGPT2 is widely accepted as an antagonist of TIE2 and destabilizes blood vessels by suppressing mural cell recruitment [15, 29], we reasoned that the abnormal mural cell association with tip ECs observed upon ANGPT2 insufficiency might be driven by increased TIE2 signaling. To test this hypothesis and investigate its broader consequences on cerebrovascular angiogenesis, we genetically deleted *Tek* (the gene encoding TIE2) in the endothelium. Endothelial *Tek* deletion was induced via intraperitoneal tamoxifen injections from postnatal days 1 to 3 (P1–P3), and cerebrovascular phenotypes were analyzed at P11–P15. Successful knockout was confirmed by co-immunostaining for TIE2 and the endothelial marker PECAM1 (**Fig. 4D**). Gross morphological evaluation revealed prominent bleeding phenotypes in the cerebellum (**Fig. 4D**), which were corroborated by H&E staining and TER119/PECAM1 immunofluorescence imaging in the cerebrum of *Tek^iKO;CDH5+^* mice (**Fig. 4E**). Furthermore, tuft-like neovascular structures were detected across multiple brain regions, including the cortex, hippocampus, and thalamus of *Tek^iKO;CDH5+^* mice at P15 (**Fig. 4F**). Suprisingly, despite TIE2’s canonical role in mural cell recruitment, endothelial TIE2 attenuation did not seem to alter perivascular cell coverage in the brain, as assessed by PDGFRβ and NG2 immunostaining (**Fig. 4G**), recapitulating our prior findings in the retina [22]. Finally, endothelial junctional analysis demonstrated that the adhesion junction molecule CDH5 displayed a fragmented, discontinuous pattern within the neovascular tufts, contrasting sharply with the linear, continuous structure observed in littermate controls (**Fig. 4H**).

### Aberrant mural cell-tip EC coupling within 48 h post-*Angpt2* deletion

To obtain direct evidence of abnormal mural cell attachment to tip ECs caused by ANGPT2 insufficiency, endothelial-specific *Angpt2* deletion was induced in mice at P4-P5, and the retinas were analyzed 48 hours later. The tip ECs in the angiogenic front of *Angpt2^iKO;CDH5+^* retinas exhibited aberrant pericyte recruitment, accompanied by a significant increase in ESM1-positive cells and a marked reduction in the total vascularized area (Fig. 5A-C). This closely mirrors the phenotype observed at P7 following postnatal endothelial ANGPT2 insufficiency. To further validate these early changes, endothelial *Angpt2* deletion was induced at P4 and the retinas were examined 24 hours later (Supplemental Fig. 3A). This earlier time point recapitulated the key abnormalities, including aberrant pericyte recruitment on tip cells, increased ESM1-positive cells, and reduced vascularized area in *Angpt2^iKO;CDH5+^* mice (Supplemental Fig. 3B-E). Beyond ANGPT2, other angiopoietin family members, including ANGPT1 and ANGPT4 also contribute to vascular development. To assess their roles, we generated a conditional inducible *Angpt1* knockout mouse model using the *Ubc-Cre^ERT2^* mouse strain (*Angpt1^Flox/−^*;*Ubc-Cre^ERT2^*, designated as *Angpt1^iKO;UBC+^*), utilizing *Angpt1^Flox/+^*; *Ubc-Cre^ERT2^* littermates as controls. Following induced gene deletion from embryonic 12.5 days (E12.5) to E14.5, vascular development was examined at E18.5. *Angpt1^iKO;UBC+^* mice displayed delayed cutaneous venous development and mis-patterned mesenteric veins (Supplemental Fig. 4A-D), consistent with previously reported venous defects in the *Tek* or *Tie1* mutant mice [22, 23]. Furthermore, immunofluorescence staining revealed significantly reduced diameters of both central and peripheral retinal veins in *Angpt4*^−/−^ mice at P21 (Supplemental Fig. 4E, F), as also reported by Elamaa et al. [24]. Collectively, these findings demonstrate that distinct angiopoietins play specialized, non-redundant roles in vascular development.

**Fig. 5.**
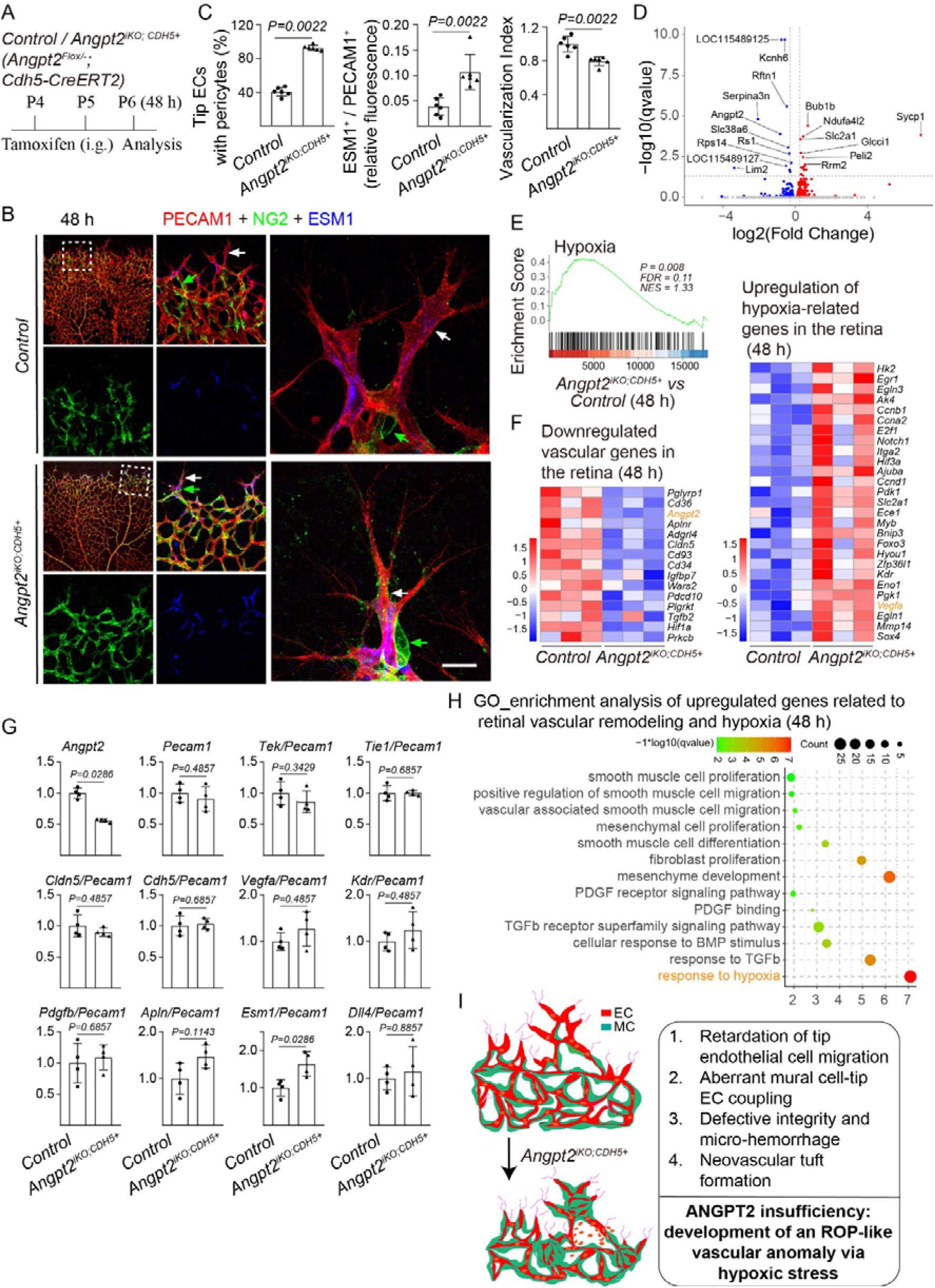
Transcriptomic alterations related to aberrant mural cell association with tip cells within 48 hours of induced endothelial *Angpt2* deletion. **A.** Tamoxifen intragastric (i.g.) administration and analysis scheme. **B**. Immunostaining for NG2 of retina from *Angpt2^iKO;CDH5+^* and littermate control mice at P6. Green arrows point to pericytes and white arrows to tip ECs. **C**. Quantification of pericyte association with tip ECs in *Angpt2^iKO;CDH5+^* and littermate control mice at P6. The ratio of ESM1 positive tip ECs with pericytes coverage to total tip ECs (Control: 40.64±5.17 %, n=6; *Angpt2^iKO;CDH5+^*: 92.76±3.15 %, n=6, *P*=0.0022). Relative fluorescence intensity of ESM1-positive ECs at the retinal angiogenic front, normalized to PECAM1 (Control: 0.036±0.0037, n=6; *Angpt2^iKO;CDH5+^*: 0.094±0.013, n=6, *P*=0.0022). The vascularization index, defined as the ratio of vascularized area to total retinal area normalized to littermate controls (Control: 1.00±0.10, n=6; *Angpt2^iKO;CDH5+^*: 0.80±0.05, n=6, *P*=0.0022). **D**. The Volcano plot of differentially regulated genes at 48 hours after *Angpt2* deletion. **E**. GSEA analysis revealed a significant increase in hypoxia-related gene expression following *Angpt2* deletion. **F.** Heatmap showing subsets of downregulated genes related to vascular formation and integrity, and upregulated gene (*P<0.05*) associated with hypoxia in retina. **G**. Quantitative expression analysis of vascular integrity genes (*Tek*, *Tie1*, *Cldn5*, *Cdh5*) and angiogenesis-related genes (*Vegfa*, *Kdr*, *Pdgfb*, *Apln*, *Esm1*, *Dll4*) in retinal tissues at 48 hours after *Angpt2* deletion. **H**. GO enrichment analysis showed that upregulated genes were associated with retinal vascular remodeling and hypoxia at 48h. **I**. Schematic illustration showing that the ANGPT2 insufficiency leads to abnormal recruitment of pericytes to tip ECs, suppression of endothelial cell sprouting and migration, and impaired vascular integrity. Scale bar: 20 μm in B.

Retinal RNA-seq 48 hours after endothelial *Angpt2* deletion revealed distinct differentially expressed genes compared to littermate controls (Fig. 5D). GSEA demonstrated significant enrichment of hypoxia-related pathways, which were elevated within 48 hours following ANGPT2 insufficiency (Fig. 5E). For instance, *Esm1*, a downstream target of the VEGF signaling, was significantly upregulated. Heatmap analysis displayed a downregulation of genes associated with vascular formation and integrity alongside an upregulation of genes linked to hypoxia (Fig. 5F), closely mirroring the findings at P7. Quantification of differentially expressed genes revealed a downward trend in vascular integrity-related genes *Tek* and *Cldn5*, while the hypoxic marker *Esm1* was significantly upregulated, and *Vegfa*, *Kdr*, *Apln*, and *Dll4* also exhibited upward trends (Fig. 5G). Furthermore, GO enrichment of the upregulated genes highlighted biological processes related to hypoxic responses as well as vascular smooth muscle cell migration and proliferation (Fig. 5H). Taken together, these findings indicate that ANGPT2 insufficiency suppresses retinal tip endothelial cell migration, thereby promoting hypoxia-triggered aberrant vascular formation (Fig. 5I).

## Discussion

ANGPT2 has historically been implicated as a key factor driving pathological angiogenesis in retinopathy of prematurity (ROP). Through analysis of single-cell RNA-seq data of the neonatal retina [27], we found that *Angpt2* transcripts were highly enriched in tip cells compared to other endothelial subpopulations, where the expression of other angiopoietins or their TIE2 receptor is minimal or absent. To further characterize the role of ANGPT2 in tip ECs, we induced endothelial-specific deletion of *Angpt2* and demonstrate that ANGPT2 insufficiency disrupts initial tip cell migration. At the molecular level, a hypoxia-driven transcriptional signature, including the upregulation of *Vegfa* and *Pdgfb*, was detected in the retinas of *Angpt2^iKO;CDH5+^* mice. This molecular shift triggered aberrant mural cell recruitment around tip ECs, defective vascular integrity, a marked increase in ESM1-positive proliferating ECs, and the formation of neovascular tufts. This complex phenotype closely phenocopies the vascular pathology characteristic of retinopathy of prematurity (ROP). Findings of this study imply that ANGPT2 serves as a pivotal regulator of tip EC invasion, and its insufficiency triggers a localized hypoxic cascade that ultimately culminates in an ROP-like vascular anomaly.

While ANGPT1 insufficiency or the loss of ANGPT4 affected developmental processes of venous morphogenesis as shown in this study and also by other researchers [21, 24], the induced endothelial *Angpt2* deletion suppressed sprouting angiogenesis. This phenotype, together with the RNA-seq data from the retinas of *Angpt2* mutants, suggests that the tip cell-derived ANGPT2 is crucial for endothelial invasion at the vascular fronts. Tip cells are a transient type of endothelial cells, defined by their anatomical position and distinct morphology with filopodia protrusions. They are highly invasive cells specialized for the guidance of growing vessels at the leading edge during vascular morphogenesis. As TIE2 is lowly expressed in these cells [22], the tip cell migration is likely to be regulated in a TIE2-independent manner. Indeed, ANGPT2 binds directly to integrins such as α5β1 and activates the downstream FAK (focal adhesion kinase) signaling for EC migration [26, 30]. Consistently, ANGPT2 has also been shown to induce lymphatic EC migration, independent of TIE receptors, via β1 integrin-mediated RhoA activation [31]. Furthermore, it has also been reported that ANGPT2 drives extracellular matrix (ECM) degradation at the vascular front via the integrin-dependent activation of matrix metalloproteinases (such as MMP-2 and MMP-9) [32]. This may, at least partially, account for the defective tip cell invasion observed in the *Angpt2^iKO;CDH5+^* mutant mice. Addtionally, it is worth pointing out that endothelial cells rely on glycolysis for ATP production, particularly for the invasive tip cells at the hypoxia zone of vascular fronts [33]. It has been shown that ANGPT2 is the downstream target of PKM2 (pyruvate kinase type M2) in endothelial cells [34]. It is therefore likely that ANGPT2 insufficiency may disrupt endothelial metabolism, leading to the retarded sprouting angiogenesis.

One key feature of tip cells at the vascular fronts is that they are free of mural cell coverage. It has been widely accepted that the angiogenic vessel growth is dependent on the synergy of VEGFA and ANGPT2 [16, 35]. VEGFA induces the expression of ANGPT2 via the activation of calcineurin-NFAT pathway and triggers its release from Weibel-Palade bodies into the extracellular matrix [36, 37]. ANGPT2 is typically regarded as an antagonist to ANGPT1 and promotes the detachment of pericytes away from the vessel wall, making vascular wall more prone to sprouting angiogenesis in response to growth cues such as VEGFA [4, 33, 38]. We showed in this study that ANGPT2 insufficiency triggers a hypoxia-driven transcriptomic reprogramming. A rapid induction of PDGFB and aberrant mural cell association with invading tip cells was observed within 48 hours post induced *Angpt2* deletion. Therefore, mural cell recruitment may be secondary to hypoxia-driven factors, arising from defective endothelial invasion.

Interestingly, we have shown pathological evidences that ANGPT2 insufficiency (*Angpt2^iKO;CDH5+^*) could drive the retinopathy of prematurity (ROP)-like vascular defects. In clinical setting, ROP is a biphasic disease initiated by hyperoxia-induced vessel obliteration, followed by a compensatory but highly aberrant hypoxic phase marked by pathological neovascularization. A hallmark of this pathology, reproducible in oxygen-induced retinopathy (OIR) animal models, is the formation of “ROP-like neovascular tufts”. These tufts represent highly disorganized clusters of endothelial cells that fail to properly invade the avascular tissue. Accumulating evidence suggests that these tufts are not merely a product of over-proliferation, but rather an outcome of suppressed or misdirected tip cell invasion. Disruption in the regulatory mechanisms governing tip cell invasion and migration often lead to severe vascular pathologies, characterized by incomplete vascularization and the formation of abnormal, leaky vascular structures. It remains to be investigated whether there are loss of function mutations with *Angpt2* in ROP patients.

In summary, our findings offer a potential reconciliation to a long-standing paradox in vascular biology regarding the role of Angiopoietin-2 (ANGPT2) in pathological ocular angiogenesis. Traditional paradigms frequently categorize ANGPT2 as a primary, direct driver of abnormal angiogenesis and neovascularization, noting its upregulation in ischemic retinopathies including ROP. However, this conventional view fails to account for why ROP-like phenotypes paradoxically develop when endothelial ANGPT2 is genetically insufficient. The expression of ANGPT2 and PDGFB is regulated by the common upstream factor HIF1α in endothelial cells [39]. Evidences from our study show that ANGPT2 insufficiency fundamentally impairs the initial phase of physiological vascularization, thereby setting off a cascade of the hypoxia-driven transcriptome signature that promotes pathological overgrowth in the retinas of neonatal mice. One striking phenotype is the increase of PDGFB upon the endothelial *Angpt2* deletion, which may directly account for the aberrant recruitment of mural cells with vascular sprouts and blocks further angiogenic growth. Therefore, rather than acting as a direct stimulator of excessive angiogenesis including neovascular tuft formation in retinas, ANGPT2 promotes endothelial tip cell migration in response to tissue ischemia for the expansion of vascular network. More in-depth studies are warranted to investigate whether germline ANGPT2 missense mutations or copy-number deletions identified in patients correlate with the susceptibility or severity of ischemic retinopathies like ROP [40], offering a foundation for novel angiomodulatory therapies.

## Methods and Materials

### Animal models

All animal experiments were performed in accordance with the institutional guidelines of Soochow and Nanjing University Animal Center (MARC-AP#YH2 / SUDA20250507A03). All the mice used in this study were housed in a SPF (specific pathogen free) animal facility with a 12/12 hours dark / light cycle, and were free to food and water access. Normal mouse diet (Suzhou Shuangshi Experimental Animal Feed Technology Co., Ltd.) and cage bedding (Suzhou Baitai Laboratory Equipment Co., Ltd.) were used. The *Angpt2^Flox/Flox^* and *Tek^Flox/Flox^* mouse line was generated as previously described [22, 41]. The mouse line with *Angpt1* knockout first allele (*Angpt1^tm1a(KOMP)Wtsi^*) was established from EUCOMM embryonic stem cells (EPD0585-1A04), in which targeting cassette is recombined downstream of exon 2 (with exon 3 floxed). To obtain the *Angpt1^Flox/Flox^*mouse line, the lacZ reporter and neo-cassette were removed using the FLPeR mice as previously described [42]. The mouse line with *Angpt4* knockout first allele (*Angpt4^tm1a(KOMP)Wtsi^*, mice homozygous for the allele labelled as *Angpt4^−/−^*) was generated from EUCOMM embryonic stem cells (EPD0761-2F08), in which targeting cassette is recombined downstream of exon 3 (with exon 4 floxed). To generate mice with ubiquitous, endothelial or hematopoietic cell specific gene deletion mouse models, we employed the *Ubc-Cre^ERT2^* [43], *Cdh5-Cre^ERT2^*[44] and *Vav-iCre* mouse lines [45]. In all the phenotype analysis, wildtype or heterozygous littermates were used as controls. For the genotyping of *Angpt2* knockout allele, the primers used were forward primer (5’- AAGCTTGCCATG TCCAAGCTC-3’) and reverse primer (5’- GAAGCCGCGGGTGCATGCAAGTGAGTGAATGTG -3’), to amplify a 626 bp fragment for the knockout allele. For the genotyping of *Angpt1* knockout allele, the primers used were forward primer (5’- TTCCAGTCAGTGCAGGTAGAGA -3’) and reverse primer (5’- CCATGATCCGAAGAGTGGAGT -3’), to amplify a 475 bp fragment for the knockout allele. For the genotyping of *Angpt4*^+/−^ allele, the primers used were forward primer (5’- TTCCAGTCAGTGCAGGTAGAGA -3’) and reverse primer (5’- CCATGATCCGAAGAGTGGAGT -3’), to amplify a 336 bp fragment for the wild-type allele and a 499 bp fragment for the knockout allele. The genetic background of all mice is on C57BL/6J or C57BL/6J /SV129. Mice were sacrificed by asphyxiation with rising concentration of carbon dioxide gas, followed by cervical dislocation and tissue collection.

### Induced gene deletion

Induction of gene deletion was performed as previously described by tamoxifen treatment [22, 46]. Briefly, newborn pups received four daily intragastric injections of tamoxifen (60 µg / day, dissolved in sunflower seed oil; Sigma-Aldrich, T5648-5G) starting at postnatal day 1 or other specified time points. For the gene deletion at embryonic stages, pregnant mice were treated by the intraperitoneal injection of tamoxifen at E12.5–14.5 (1 mg / day). The genotypes of the *Angpt2* knockout and control mice are as follows: *Angpt2^Flox/−^*; *Ubc-Cre^ERT2^*labelled as *Angpt2^iKO;UBC+^*, *Angpt2^Flox/−^*;*Cdh5-Cre^ERT2^* labelled as *Angpt2^iKO;CDH5+^*, *Angpt2^Flox/Flox^*; *Vav-iCre* labelled as *Angpt2^KO;VAV+ (Vav-iCre)^* and littermate control mice (labelled as *control*) are heterozygous for the floxed allele or wild type mice. The genotypes of the *Angpt1* knockout and control mice are as follows: *Angpt1^Flox/−^*; *Ubc-Cre^ERT2^* labelled as *Angpt1^iKO;UBC+^* and *Angpt1^Flox/+^*; *Ubc-Cre^ERT2^* or *Angpt1^Flox/+^* used as control. The genotypes of the *Tek* knockout and control mice are as follows: *Tek^Flox/−^*;*Cdh5-Cre^ERT2^* labelled as *Tek^iKO;CDH5+^* and *Tek^Flox/+^*;*Cdh5-Cre^ERT2^* or *Tek^Flox/+^* used as control. Tissues were collected for the analysis at embryonic day 18.5 and postnatal day 5-21. The retina dissection and analysis were performed according to the protocol by Pitulescu et al [47]. Retinal vascularization index was quantified as the ratio of vascularized area to total retinal area as previously published [22].

### Single cell RNA-seq data analysis

The single-cell RNA sequencing data of postnatal day 6 (P6) mouse retina from Zarkada et al. were re-analyzed using the Seurat package (v5.2.1) [27]. All parameters and criteria were applied as described in the original study to ensure direct comparability. Endothelial cells (ECs) were initially identified based on the expression of pan-EC markers *Pecam1* and *Cldn5*. And then re-clustered by 2000 highly variable genes, principal components dims = 12 and resolution = 0.6 into five major EC subtypes: venEC (venous EC), artEC (artery EC), capEC(capillary EC), tip EC (tipEC), proEC (proliferated EC). The major cell types were annotated based on established marker genes: venEC (*Nr2f2*, *Emcn*), artEC (*Dll4*, *Bmx*), capEC (*Mfsd2a*, *Slc22a8*), tipEC (*Flt4*, *Kcne3*) and proEC (*Mki67*, *Birc5*). The “other” cell cluster exhibits low expression of endothelial cell markers, suggesting it may represent contamination by other cell types or doublets. Unsupervised clustering of cells was performed by RunUMAP, FindNeighbors and FindClusters, and the expression of marker genes was visualized with the FeaturePlot function.

### Bulk RNA sequencing analysis

For the bulk RNA-seq analysis of retinal tissues, eyeballs were harvested from the *Angpt2* mutants and littermate *control* mice. The retinas were dissected and snap-frozen in the liquid nitrogen. Total RNA was isolated from the retinas using Trizol (Invitrogen, 15596026CN) following the manufacturer’s instructions. The procedures for the RNA-seq analysis were performed as previously reported [23]. Briefly, sequencing was performed on DNBSEQ platform with PE150 (read length, BGI-Shenzhen, China), and the sequencing data were processed according to the predefined analysis pipeline. The differential expression analysis was conducted using the R package DESeq2 (v 1.42.1). Functional enrichment analysis was performed using clusterProfiler (v4.10.1) and org.Mm.eg.db (v 3.18.0). The hallmark gene sets used in the GSEA (Gene Set Enrichment Analysis), including the hallmarks for retinal structural ECs and tip ECs, are mainly defined according to the single cell transcriptome analysis of murine retinas, the molecular signatures database (mh.all.v2024.1.Mm.symbols) and the related GO terms [23, 48–50].

### Western blot analysis

Lung tissues were collected and lysed in NP-40 lysis buffer (Beyotime P0013F) supplemented with protease inhibitor cocktail (complete Mini, Roche 04693124001), phosphatase inhibitor cocktail (PhosSTOP, Roche 04906837001), and 1 mM PMSF (Bio Basic, PB0425). Protein concentration was measured using the BCA protein assay kit (PIERCE, 23227), and equal amounts of protein were analyzed. Following transfer from gels to PVDF membranes (IPVH00010, Millipore) and antibody incubation, images were acquired by the chemiluminescent detection method (NEL105001EA, PerkinElmer) using X-ray film (XBT, Carestream, 6535876). Protein markers were manually annotated on the films by overlapped the PVDF membranes with a prestained protein ladder (ThermoFisher Scientific, 26616). The blots were washed and re-probed with antibodies for beta-Actin as loading controls. The following antibodies were used in this study, including sheep anti-ANGPT2 (R&D Systems, AF7186, 1:500), mouse anti-beta-Actin (Santa Cruz, sc-47778 HRP, 1:10000) and rabbit-anti-sheep IgG, HRP Conjugated (Proteintech, SA00001-16, 1:5000).

### Immunostaining

For the whole-mount immunostaining, tissues including retina, intestinal villi, skin and mesentery were harvested and processed as previously described [46, 51]. Briefly, samples were fixed in 4% paraformaldehyde (PFA, Sigma-Aldrich, 158127) for 2 hours, and blocked with 3% (w/v) skim milk (Valio, skimmed milk powder instant) in PBS containing 0.3% Triton X-100 (VETEC, V900502), and then incubated the primary antibodies at 4℃ overnight. For immunofluorescence staining of retinal and cerebral sections, tissues were fixed in 4% PFA for 2 hours, dehydrated in 20% sucrose solution at 4°C overnight, and embedded in tissue freezing medium. The following antibodies were used: rat anti-PECAM1 (BD Biosciences, 553370, 1:500), goat anti-DLL4 (R&D Systems, AF1389, 1:500), goat anti-ESM1 (R&D Systems, AF1999, 1:500), rabbit anti-NG2 (Merck Millipore, AB5320, 1:500), rabbit anti-CLDN5 (Invitrogen, 34-1600, 1:500), goat anti-CDH5 (R&D Systems, AF1002, 1:500), rat anti-TER119 (eBioscience, 14-5921, 1:500), and eFluor 660-conjugated mouse anti-αSMA (eBioscience, 50-9760-82, 1:500), goat anti-TIE2 (R&D Systems, AF762, 1:500) and goat anti-PDGFRβ (R&D Systems, AF1042, 1:500). The secondary antibodies used were: Alexa Fluor 488-conjugated donkey anti-rat IgG (Invitrogen, A-21208, 1:500), Alexa Fluor 488-conjugated donkey anti-rabbit IgG (Invitrogen, A-21206, 1:500), Alexa Fluor 568-conjugated donkey anti-rat IgG (Invitrogen, A78946, 1:500), Cy3-conjugated donkey anti-goat IgG (Jackson ImmunoResearch, 705-165-147), Cy5-conjugated donkey anti-goat IgG (Jackson ImmunoResearch, 705-175-147), and Cy5-conjugated donkey anti-rat IgG (Jackson ImmunoResearch, 712-175-153). Fluorescently labeled samples were mounted with 50% glycerol (Sigma-Aldrich, G5516) containing DAPI (Beyotime, C1002) and analyzed with a confocal microscope (Olympus Fluoview 1000 or 3000). Imaging parameters were kept consistent across all confocal analyses for comparative purposes.

### Statistical analysis

For the statistical analysis of 2-group comparison, the unpaired t test was performed with Welch correction if data passed the D’Agostino-Pearson normality test, or the unpaired nonparametric Mann-Whitney U test was applied using GraphPad Prism 8. Data are expressed as mean±SD. All statistical tests were 2-sided.

## Data availability

The raw and processed RNA-seq data generated in this study are to be deposited in the NCBI Gene Expression Omnibus (GEO) database and made publicly available upon publication.

## Authorship Contributions

Z.S., K.D., T.L conducted the experiments, acquired and analyzed data, and wrote the manuscript. J.Z., X.S., X.J., X.L., X.C., B.X., P.L. provided technical supports and critical comments on this study. Y.H. designed, organized, supervised the project, analyzed data and wrote the manuscript.

## Acknowledgement and funding support

We thank staff in the Animal facility of Soochow University for technical assistance. This work was supported by grants from the National Natural Science Foundation of China (82470518), Cyrus Tang Foundation (CTJC25001), China Postdoctoral Science Foundation (CPSF, 2024M762300), the National Natural Science Foundation of China (82401544), the National Key R&D Program of China (2021YFA0805000), the Natural Science Foundation of Jiangsu Province (BK20240786), and the Priority Academic Program Development of Jiangsu Higher Education Institutions.

## Reference

1. Lee, H.W., et al., Role of Venous Endothelial Cells in Developmental and Pathologic Angiogenesis. Circulation, 2021. 144(16): p. 1308–1322.

2. Gerhardt, H., et al., VEGF guides angiogenic sprouting utilizing endothelial tip cell filopodia. J Cell Biol, 2003. 161(6): p. 1163–77.

3. Blanco, R. and H. Gerhardt, VEGF and Notch in tip and stalk cell selection. Cold Spring Harb Perspect Med, 2013. 3(1): p. a006569.

4. Hellstrom, M., et al., Dll4 signalling through Notch1 regulates formation of tip cells during angiogenesis. Nature, 2007. 445(7129): p. 776–80.

5. Rocha, S.F., et al., Esm1 modulates endothelial tip cell behavior and vascular permeability by enhancing VEGF bioavailability. Circ Res, 2014. 115(6): p. 581–90.

6. Pitulescu, M.E., et al., Dll4 and Notch signalling couples sprouting angiogenesis and artery formation. Nat Cell Biol, 2017. 19(8): p. 915–927.

7. Gaengel, K., et al., Endothelial-mural cell signaling in vascular development and angiogenesis. Arterioscler Thromb Vasc Biol, 2009. 29(5): p. 630–8.

8. Siekmann, A.F., Biology of vascular mural cells. Development, 2023. 150(16).

9. Park, D.Y., et al., Plastic roles of pericytes in the blood-retinal barrier. Nat Commun, 2017. 8: p. 15296.

10. Hellstrom, M., et al., Role of PDGF-B and PDGFR-beta in recruitment of vascular smooth muscle cells and pericytes during embryonic blood vessel formation in the mouse. Development, 1999. 126(14): p. 3047–55.

11. Lindahl, P., et al., Pericyte loss and microaneurysm formation in PDGF-B-deficient mice. Science, 1997. 277(5323): p. 242–5.

12. ten Dijke, P. and H.M. Arthur, Extracellular control of TGFbeta signalling in vascular development and disease. Nat Rev Mol Cell Biol, 2007. 8(11): p. 857–69.

13. Yu, J., et al., Endothelial nitric oxide synthase is critical for ischemic remodeling, mural cell recruitment, and blood flow reserve. Proc Natl Acad Sci U S A, 2005. 102(31): p. 10999–1004.

14. Ando, K., et al., Peri-arterial specification of vascular mural cells from naive mesenchyme requires Notch signaling. Development, 2019. 146(2).

15. Augustin, H.G., et al., Control of vascular morphogenesis and homeostasis through the angiopoietin-Tie system. Nat Rev Mol Cell Biol, 2009. 10(3): p. 165–77.

16. Saharinen, P., L. Eklund, and K. Alitalo, Therapeutic targeting of the angiopoietin-TIE pathway. Nat Rev Drug Discov, 2017. 16(9): p. 635–661.

17. Li, J.J., et al., Thrombin induces the release of angiopoietin-1 from platelets. Thromb Haemost, 2001. 85(2): p. 204–6.

18. Sundberg, C., et al., Stable expression of angiopoietin-1 and other markers by cultured pericytes: phenotypic similarities to a subpopulation of cells in maturing vessels during later stages of angiogenesis in vivo. Lab Invest, 2002. 82(4): p. 387–401.

19. Saharinen, P., et al., Angiopoietins assemble distinct Tie2 signalling complexes in endothelial cell-cell and cell-matrix contacts. Nat Cell Biol, 2008. 10(5): p. 527–37.

20. Fukuhara, S., et al., Differential function of Tie2 at cell-cell contacts and cell-substratum contacts regulated by angiopoietin-1. Nat Cell Biol, 2008. 10(5): p. 513–26.

21. Arita, Y., et al., Myocardium-derived angiopoietin-1 is essential for coronary vein formation in the developing heart. Nat Commun, 2014. 5: p. 4552.

22. Chu, M., et al., Angiopoietin receptor Tie2 is required for vein specification and maintenance via regulating COUP-TFII. Elife, 2016. 5:e21032.

23. Cao, X., et al., Endothelial TIE1 Restricts Angiogenic Sprouting to Coordinate Vein Assembly in Synergy With Its Homologue TIE2. Arterioscler Thromb Vasc Biol, 2023. 43(8): p. e323–e338.

24. Elamaa, H., et al., Angiopoietin-4-dependent venous maturation and fluid drainage in the peripheral retina. Elife, 2018. 7.

25. Kim, M., et al., Opposing actions of angiopoietin-2 on Tie2 signaling and FOXO1 activation. J Clin Invest, 2016. 126(9): p. 3511–25.

26. Felcht, M., et al., Angiopoietin-2 differentially regulates angiogenesis through TIE2 and integrin signaling. J Clin Invest, 2012. 122(6): p. 1991–2005.

27. Zarkada, G., et al., Specialized endothelial tip cells guide neuroretina vascularization and blood-retina-barrier formation. Dev Cell, 2021. 56(15): p. 2237–2251 e6.

28. Oh, H., et al., Hypoxia and vascular endothelial growth factor selectively up-regulate angiopoietin-2 in bovine microvascular endothelial cells. J Biol Chem, 1999. 274(22): p. 15732–9.

29. Kraft, M., et al., Angiopoietin-TIE2 feedforward circuit promotes PIK3CA-driven venous malformations. Nat Cardiovasc Res, 2025. 4(7): p. 801–820.

30. Akwii, R.G., et al., Role of Angiopoietin-2 in Vascular Physiology and Pathophysiology. Cells, 2019. 8(5).

31. Akwii, R.G., et al., Angiopoietin-2-induced lymphatic endothelial cell migration drives lymphangiogenesis via the beta1 integrin-RhoA-formin axis. Angiogenesis, 2022. 25(3): p. 373–396.

32. Hu, B., et al., Angiopoietin 2 induces glioma cell invasion by stimulating matrix metalloprotease 2 expression through the alphavbeta1 integrin and focal adhesion kinase signaling pathway. Cancer Res, 2006. 66(2): p. 775–83.

33. De Bock, K., et al., Role of PFKFB3-driven glycolysis in vessel sprouting. Cell, 2013. 154(3): p. 651–63.

34. Ge, Q., et al., PKM2 regulates angiogenic activation via ANGPT2 in endothelial cells. Sci Rep, 2025. 15(1): p. 31448.

35. Asahara, T., et al., Tie2 receptor ligands, angiopoietin-1 and angiopoietin-2, modulate VEGF-induced postnatal neovascularization. Circ Res, 1998. 83(3): p. 233–40.

36. Minami, T., et al., The calcineurin-NFAT-angiopoietin-2 signaling axis in lung endothelium is critical for the establishment of lung metastases. Cell Rep, 2013. 4(4): p. 709–23.

37. Fiedler, U., et al., The Tie-2 ligand angiopoietin-2 is stored in and rapidly released upon stimulation from endothelial cell Weibel-Palade bodies. Blood, 2004. 103(11): p. 4150–6.

38. Maisonpierre, P.C., et al., Angiopoietin-2, a natural antagonist for Tie2 that disrupts in vivo angiogenesis. Science, 1997. 277(5322): p. 55–60.

39. Kelly, B.D., et al., Cell type-specific regulation of angiogenic growth factor gene expression and induction of angiogenesis in nonischemic tissue by a constitutively active form of hypoxia-inducible factor 1. Circ Res, 2003. 93(11): p. 1074–81.

40. Leppanen, V.M., et al., Characterization of ANGPT2 mutations associated with primary lymphedema. Sci Transl Med, 2020. 12(560).

41. Shen, B., et al., Genetic dissection of tie pathway in mouse lymphatic maturation and valve development. Arterioscler Thromb Vasc Biol, 2014. 34(6): p. 1221–30.

42. Farley, F.W., et al., Widespread recombinase expression using FLPeR (flipper) mice. Genesis, 2000. 28(3-4): p. 106–10.

43. Ruzankina, Y., et al., Deletion of the developmentally essential gene ATR in adult mice leads to age-related phenotypes and stem cell loss. Cell Stem Cell, 2007. 1(1): p. 113–26.

44. Okabe, K., et al., Neurons limit angiogenesis by titrating VEGF in retina. Cell, 2014. 159(3): p. 584–96.

45. de Boer, J., et al., Transgenic mice with hematopoietic and lymphoid specific expression of Cre. Eur J Immunol, 2003. 33(2): p. 314–25.

46. Cao, X., et al., A Genetically Engineered Mouse Model of Venous Anomaly and Retinal Angioma-like Vascular Malformation. Bio Protoc, 2021. 11(15): p. e4117.

47. Pitulescu, M.E., et al., Inducible gene targeting in the neonatal vasculature and analysis of retinal angiogenesis in mice. Nat Protoc, 2010. 5(9): p. 1518–34.

48. Kalucka, J., et al., Single-Cell Transcriptome Atlas of Murine Endothelial Cells. Cell, 2020. 180(4): p. 764–779 e20.

49. Liberzon, A., et al., The Molecular Signatures Database (MSigDB) hallmark gene set collection. Cell Syst, 2015. 1(6): p. 417–425.

50. Macosko, E.Z., et al., Highly Parallel Genome-wide Expression Profiling of Individual Cells Using Nanoliter Droplets. Cell, 2015. 161(5): p. 1202–1214.

51. Zhang, L., et al., VEGFR-3 ligand-binding and kinase activity are required for lymphangiogenesis but not for angiogenesis. Cell Res, 2010. 20(12): p. 1319–31.

