## Supplemental Fig. 1-4 for "Endothelial ANGPT2 insufficiency impairs retinal vascularization with ROP-like neovascular tufts"

### Supplemental figures and figure legends

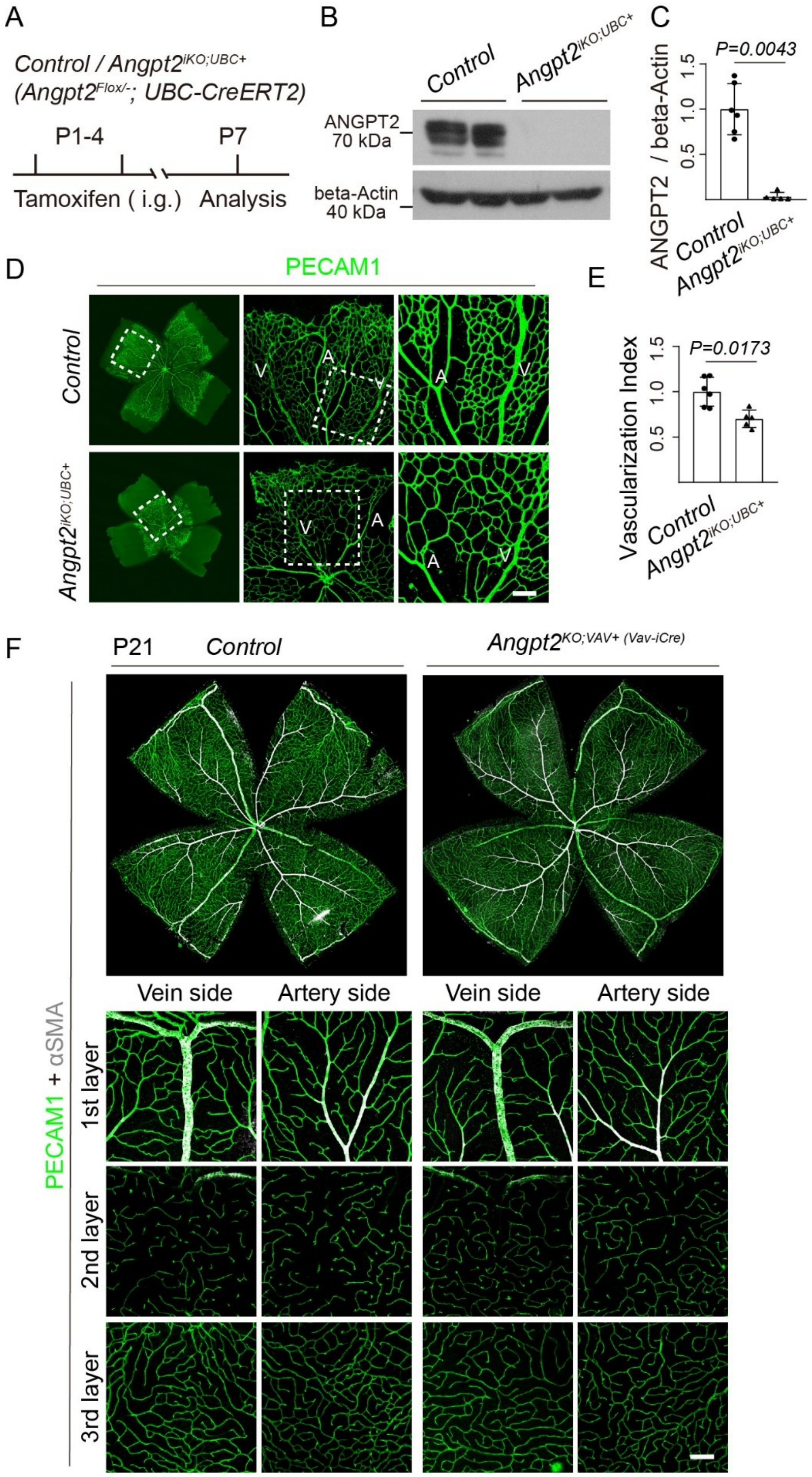

**Supplemental Fig. 1 Analysis of retinal vascular morphogenesis after the induced *Angpt2* deletion ubiquitously or in hematopoietic cells. A**. Tamoxifen intragastric (i.g.) administration and analysis scheme. **B**. Deletion efficiency of ANGPT2 was assessed by the Western blotting analysis of lung tissues from *Angpt2*$^{iKO;UBC+}$ and control mice at P7. **C**. Quantification of ANGPT2 protein levels normalized to beta-actin in lungs of *Angpt2*$^{iKO;UBC+}$ and littermate control mice at P7 (ANGPT2/beta-actin: Control: 1.00±0.28, n=6; *Angpt2*$^{iKO;UBC+}$: 0.03±0.05, n=5, *P*=0.0043). **D**. Retinal blood vessel analysis of *Angpt2*$^{iKO;UBC+}$ and control mice at P7. **E**. Quantification of the vascularization index, defined as the ratio of vascularized area to total retinal area normalized to littermate controls at P7 (Control: 1.00±0.16, n=6; *Angpt2*$^{iKO;UBC+}$: 0.70±0.10, n=5, *P*=0.017). **F**. Analysis of the three layers of retinal blood vessels in *Angpt2*$^{KO;vav+ (Vav-iCre)}$ and control mice by whole-mount immunostaining at P21. Scale bar: 100 μm in D and F.

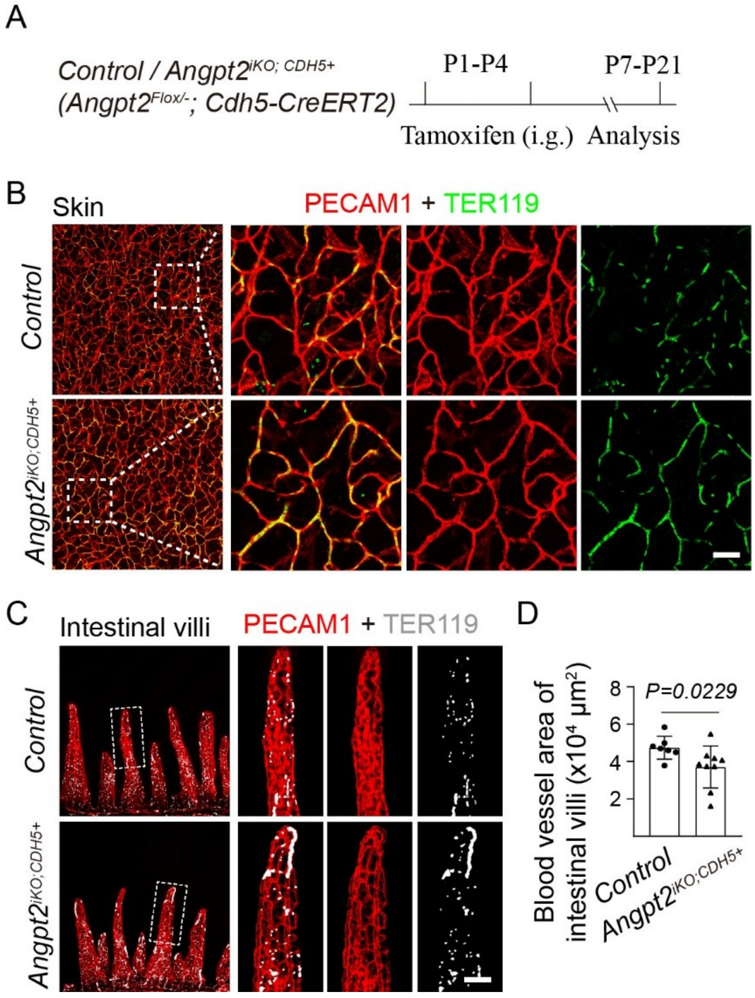


**Supplemental Fig. 2 Differential requirement of ANGPT2 in angiogenesis of skin and intestinal villi. A**. Tamoxifen intragastric (i.g.) administration and analysis scheme.

**B**. Comparisons of vascular density and integrity in control and *Angpt2*$^{iKO;CDH5+}$ mice by abdominal skin whole-mount staining at P7. **C**. Comparisons of blood vascular area and integrity in the duodenum of control and *Angpt2*$^{iKO;CDH5+}$ mice by whole-mount staining at P21. **D**. Quantification of blood vascular area in the duodenum of control and *Angpt2*$^{iKO;CDH5+}$ mice at P21 (×$10^4$ μm$^2$, Control: 4.74 ± 0.61, n=7; *Angpt2*$^{iKO;CDH5+}$: 3.71±1.13, n=9; P=0.0229). Scale bar: 50 μm in B-C.

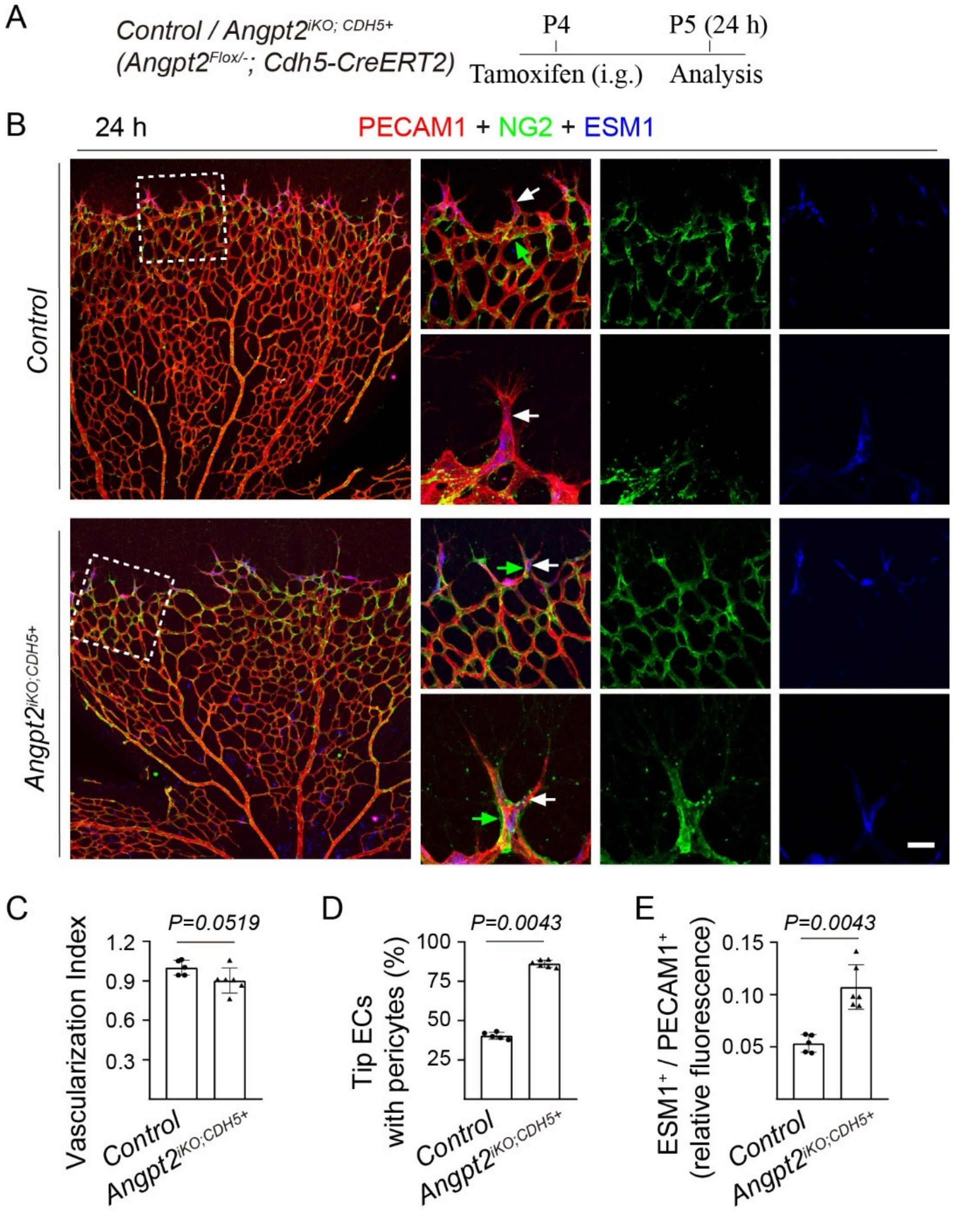


**Supplemental Fig. 3 Abnormal recruitment of mural cells within 24 hours of induced endothelial *Angpt2* deletion. A**. Tamoxifen intragastric (i.g.) administration and analysis scheme. **B**. Immunostaining for NG2 in retina from *Angpt2*$^{iKO;CDH5+}$ and littermate control mice at P5. White arrows point to tip ECs and green arrows to pericytes. **C**. Quantification of the vascularization index, defined as the ratio of vascularized area to total retinal area normalized to littermate controls (Control: 1.00 ± 0.057, n=5; *Angpt2*$^{iKO;CDH5+}$: 0.90 ± 0.095, n=6, *P*=0.0519). **D**. Analysis of pericyte association with tip ECs in *Angpt2*$^{iKO;CDH5+}$ and control mice at P5. The ratio of ESM1 positive tip ECs with pericytes coverage to total tip ECs (Control: 40.38 ± 2.16%, n=5; *Angpt2*$^{iKO;CDH5+}$: 86.21 ± 2.34%, n=6, *P*=0.0043). **E**. Relative fluorescence intensity of ESM1-positive ECs at the retinal angiogenic front, normalized to PECAM1 (Control: 0.053 ± 0.0085, n=6; *Angpt2*$^{iKO;CDH5+}$: 0.11 ± 0.021, n=6, *P*=0.0043). Scale bar: 20 μm in B.

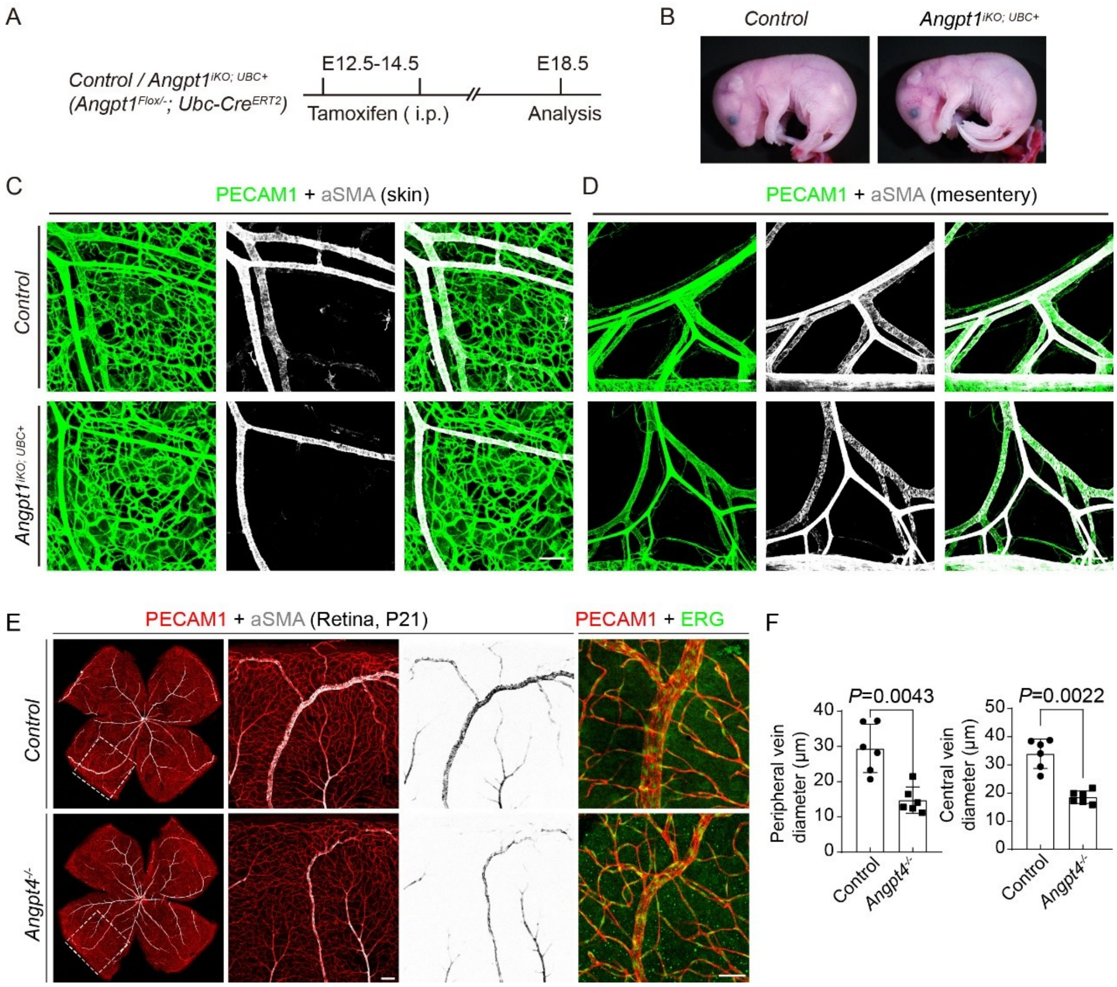


**Supplemental Fig. 4 Requirement of ANGPT1 and ANGPT4 in the regulation of vein development. A**. Tamoxifen intraperitoneal (i.p.) administration and analysis scheme. **B**. Morphological appearance of *Angpt1*$^{iKO;UBC+}$ mice and their littermate controls. **C-D**. Immunostaining for PECAM1 (green) and αSMA (gray) of skin (C) and mesentery (D) from *Angpt1*$^{iKO;UBC+}$ mice and littermate controls at E18.5. **E**. Immunostaining for PECAM1 (red), αSMA (gray) and ERG (green) in retina of *Angpt4*$^{-/-}$ and littermate control mice at P21. **F**. Quantification of retinal vein diameters in *Angpt4*$^{-/-}$ and littermate control mice at P21 (Peripheral vein diameter, Control: 29.40 ±

6.86, n=6; *Angpt4*$^{-/-}$: 14.73 ± 3.75, n=6, *P*=0.0043; Central vein diameter, Control: 33.96 ± 5.23, n=6; *Angpt4*$^{-/-}$: 18.55 ± 2.22, n=6, *P*=0.0022). Scale bar: 100 μm in C-E.
